# Dietary landscapes shape genotype- and sex-specific responses to insecticides

**DOI:** 10.64898/2026.03.02.708330

**Authors:** A. Nogueira Alves, B.J. Houston, Y.T. Yang, N. Wedell

**Author notes:** Corresponding author: André Nogueira Alves.

## Abstract

Insecticide resistance is typically studied as a response to chemical toxicity, yet in natural and agricultural systems insecticides are embedded within food resources. How resistance alleles interact with nutritional environments to shape fitness remains largely unknown. Here we combine nutritional geometry approaches to test how variation at a major resistance locus, *Cyp6g1*, modifies sex-specific reproductive performance across dietary landscapes in *Drosophila melanogaster*. We show that a resistance allele does more than increase survival: it profoundly reshapes reproductive allocation in both sexes. Resistant females exhibited up to a two-fold increase in ovariole number, with benefits amplified in protein-rich, high-calorie diets. In contrast, resistant males displayed increased testis size but reduced seminal vesicle and accessory gland size, revealing sex-specific trade-offs. Critically, contaminating doses of imidacloprid shifted nutritional optima according to genotype, in some cases enhancing reproduction in susceptible flies, consistent with diet-dependent hormesis. Thus, resistance, nutrient availability and toxin exposure jointly determine fitness outcomes. Our findings demonstrate that resistance evolution is embedded within dietary landscapes rather than driven by toxicity alone, highlighting the need to integrate nutritional ecology into predictions of resistance dynamics in human-modified environments.

## Introduction

The foods organisms exploit plays a pivotal role in defining their nutritional niche, geographic distribution and abundance (Hutchinson, 1957). Nutrition is a central component of organismal performance and manipulating the quality and quantity of diet shapes life-history traits such as longevity, and reproduction (Fontana et al., 2010; Mirth et al., 2019; Späth et al., 2020). Diet also has a vital role in immune response across taxa (Childs et al., 2019), as well as being responsible for mitigating stress to various environmental factors such as cold, desiccation, and the presence of toxins (LeBlanc, 1957; McCracken et al., 2020; Petschenka & Agrawal, 2015; van Thiel et al., 2022). Furthermore, dietary responses are not generalised amongst individuals but instead emerge from interactions between the environment and genotypes, with sex being a major factor that determines the dietary requirements for optimal fitness (Reed et al., 2010; Reddiex et al., 2013). Male and female organisms across taxa have different reproductive optima and produce gametes with different nutritional and energy requirements, and therefore a diet that is optimal for one sex, may not be as optimal for the other (Andrews et al., 2024; Maklakov et al., 2008).

To colonise new environments, animals must adapt to the nutritional, physical, and chemical properties of the food (Machovsky-Capuska et al., 2016). This adaptation often involves obtaining the ability to exploit foods that other organisms would not utilize due to its physical properties or, more frequently, due to their toxic, chemical compounds (R’kha et al., 1991; Appel et al., 1992). Classic examples include *Drosophila sechellia* and *Manduca sexta*, which have evolved tolerance to toxic secondary metabolites present in their host plants, *Morinda* and *Nicotiana*, respectively (R’kha et al., 1991; Appel et al., 1992). In these systems, resistance to dietary toxins is inseparable from nutritional adaptation, as tolerance enables access to food resources that underpin growth, reproduction, and population persistence. These examples illustrate that resistance traits do not evolve in isolation, but rather as components of broader nutritional strategies.

A human-mediated analogue is the case of agricultural systems. Not only do crops represent nutritional opportunities for insects that are vastly available and highly nutritious, but the use of insecticides in the last century has provided a novel toxic property of this new food source. In this context, insecticide resistance can be viewed as an adaptation that enables continued access to an otherwise favourable nutritional niche. One of the best-known examples of insecticide resistance involves *Cyp6g1*, a P450 gene involved in insect metabolism that has been recruited via novel mechanisms to detoxify DDT and several other xenobiotics (Daborn et al., 2002; ffrench-Constant, 2013). Genetic variation at this locus has been responsible for conferring insecticide resistance in several *Drosophila* species through similar mechanisms (Bozler, 2013; Schmidt et al., 2010). *Cyp6g1* has evolved several new allelic variants in *D. melanogaster* in the last 50 years as a response to the increased use of insecticides, with each new allelic variant conferring increased resistance to higher doses of insecticides (Schmidt et al., 2010). Importantly, changes in insecticide resistance provided by this locus have also been shown to incur differences in reproductive success for both sexes in the fruitfly *D. melanogaster* (Rostant et al 2015), with resistant females producing more offspring, but resistant males mating less frequently than their counterparts (McCart et al., 2005; Smith et al 2011; Rostant et al 2015). These findings suggest that resistance evolution reshapes life-history trade-offs differently in males and females, raising the possibility that nutritional context may further modulate these sex-specific outcomes.

Despite extensive research into the molecular mechanism(s) behind the function of this gene, there is a gap in knowledge regarding the added effect(s) of ecological factors. Little is known about how nutrition of pest insects impacts each sex independently, how resistance to insecticides further interacts with each sex, and most importantly, how the presence of insecticides might alter each response. The effects of insecticides on insect fitness have been extensively studied (Gul et al., 2023; Kliot & Ghanim, 2012), however, this research has largely centred on maximising toxicity and mortality under simplified conditions and treating insecticide exposure as an isolated variable rather than part of an integrated nutritional environment. Nevertheless, because insecticides do not present themselves as isolated chemicals and are always integrated into the dietary niche, insecticide resistance must involve a concurrent adaptation to the novel nutritional niche that harbours the insecticide. Therefore, whatever nutrients insects consume, they will ingest insecticides concurrently, whether that may be in crops where insecticides are applied, or in the outbounds of crops, where contaminating doses of insecticides are present (Christen et al., 2019; Knoepp et al., 2012). Although some studies have examined how dietary context modulates the effects of naturally occurring toxins (Hägele & Rowell-Rahier, 1999; Singer et al., 2002), there is a striking absence of research addressing how insecticides interact with diet, or how the same insecticide may exert different fitness consequences across distinct nutritional environments. Comparing the impact of diet composition and insecticide presence on life history traits between susceptible and resistant individuals can further our understanding of adaptation to new nutritional niches or even persistence of insect populations in the niches they occupy.

*Drosophila melanogaster* offers a powerful system for a holistic analysis of insecticide resistance. Not only does it represent an exemplary case study of *Cyp6g1* allelic diversification, but the role of nutrition on fitness has been extensively researched. Remarkably, the interaction between nutrition, resistance expression and exposure to insecticides has yet to be understood. Previous research has shown that reproduction is sensitive to larval nutrition, which is one of the biggest determinants of fitness (Wallin et al., 1992; Behmer et al., 2001; Lee et al., 2008). In female flies, ovariole number is the key trait that determines the upper limit of egg production (David, 1970). This trait has been shown to be maximised in high protein diets, and it is finely tuned by this nutrient (Rodrigues et al., 2015, Mirth et al., 2019; Alves et al., 2022). Male fertility is more complex and is not determined by one single trait. Male reproductive tracts consist of three main structures: testes, seminal vesicles (SVs), and accessory glands (AGs), all of which have essential roles in producing and maturing spermatozoa (Houston et al., 2025). Testes are responsible for producing spermatozoa (Siddall & Hime, 2017), while seminal vesicles store and mature these spermatozoa (Yuan et al., 2019). When ready to reproduce, spermatozoa are mixed with peptides produced by the accessory glands that both activate these cells as well as interact with the female’s reproductive and other organ systems to produce both physiological and behavioural changes (Wilson et al., 2017, Cohen & Wolfner, 2018; Wigby & Chapman, 2005). Due to its complexity, the role of nutrition on male fertility has not been as extensively studied as other traits. Protein levels influence male remating rates and seminal vesicle length (Fricke et al., 2008; Amitin & Pitnick 2007), but it remains unknown how different levels of carbohydrates and protein impact the size of the male reproductive tract.

Here, we make use of a susceptible and a resistant strain of *D. melanogaster* with the same genetic background (Canton-S) to understand how: 1) differences in their nutritional niche, 2) the presence or absence of either a resistance allele, 3) exposure to insecticides, or 4) both resistance alleles and insecticides, affect the response of reproductive traits to the macronutrient composition of the larval diet. We use the Nutritional Geometry Framework (Simpson & Raubenheimer, 1993; Simpson & Raubenheimer, 1999; Silva-Soares et al., 2019; Chakraborty et al., 2020) to test the hypothesis that resistant *D. melanogaster* larvae perform better on diets containing insecticides when compared with their susceptible counterparts. We know that resistance alleles at the *Cyp6g1* locus incurs a fecundity boost for female flies, however, we have no information if these alleles will also incur sex specific changes in response to nutrition. Potentially, a boost in female fertility due to a resistant allele may be further exacerbated in ideal dietary niches such as high protein diets. However, this may not hold true for male fertility, as we know males have different dietary requirements, and so a resistant allele at the *Cyp6g1* locus might only be beneficial in a male-optimal diet such as a high carbohydrate diet (Camus et al., 2017). Our approach uses artificial diets varying the quantities of two nutrients (protein and carbohydrates) to determine how various reproductive traits map to the nutritional space. It has previously been applied not only to a variety of *Drosophila* contexts, but also to many organisms across taxa (Silva-Soares et al., 2019; Chakraborty et al., 2020; Fanson et al., 2012; Raubenheimer & Simpson, 2003). Despite extensive work on nutrition, insecticide resistance, and sex-specific fitness, no studies have quantified how insecticide exposure interacts with nutritional environments to shape genotype- and sex-specific fitness outcomes. Building on evidence that *Cyp6g1*-mediated resistance enhances female fecundity and that males and females have distinct nutritional optima, we predict that resistance-associated fitness benefits will be amplified in female-optimal diets but may be constrained or reversed in males. Exploring these interactions will reveal how adaptation to insecticides is modulated by diet and sex, and how human-made chemical environments are reshaping evolutionary trajectories and population persistence in the wild.

## Methods

### Fly stocks

We used the insecticide susceptible strain Canton S (*Cyp6g1* – M allele; BDSC # 64349), and a resistant strain, which was generated by backcrossing Hikone-R (*Cypg61* – BA allele; BDSC # 4267) into a Canton S strain for 8 generations, whilst continuously selecting for the resistant allele (Rostant et al., 2015). This process has produced reliable resistant flies with extensive proof of their survivability on high doses of insecticides (Martelli et al., 2015; Nogueira Alves et al., 2026). Flies were reared at 25 ° C with a 12h:12h day:night light cycle, on sugar-yeast medium, consisting of 100 g autolysed yeast powder, 50 g sucrose, 10 g agar, 30 mL Nipagin, and 3 mL of propionic acid per liter of water (Grandison et al., 2009).

### Nutritional Geometry Framework

Twenty-five diets were used in this study, which encompasses a vast nutritional space. These diets were one of five protein and carbohydrate (P:C) ratios; 1:8, 1:4, 1:2, 1:1, 1.5:1, made by changing the quantities of autolysed yeast, and sucrose. Each ratio was prepared at 5 caloric concentrations: 1600 kcal, 1250 kcal, 900 kcal, 550 kcal, and 200 kcal. For each one of these diet combinations, a solution of either dimethyl sulfoxide (DMSO) or imidacloprid diluted in DMSO (to a concentration of 0.1 ppm/mL) was added to the food to measure the effect of low doses of insecticides on larval diet. This resulted in a total of fifty diets. Although levels of insecticide residues in the field vary depending on the formulation, application method, and environmental conditions, the final concentration of 0.1 ppm/mL is known to be a contaminating dose found in nature (Boul et al., 1994).

To obtain focal flies for the experiment, parental flies from each of the two strains were acclimated to egg laying chambers containing apple-juice plates smeared with yeast paste for 24 hours. Plates were changed every day and housed at 25 °C with a 12h:12h day:night cycle. Freshly hatched L1 larvae were collected from these plates, and 30 larvae were transferred from each strain into vials containing each of the experimental diets. Seven replicate vials for each strain, and for each of the 25 diets, were placed at 25 °C with a 12h:12h day:night cycle, where they were left to develop from egg to adult eclosion. At 3 days of age (Zamore & Ma, 2011), male and female flies were submerged in 4% paraformaldehyde for 30 min then stored at 4° C until use.

### Ovariole number and male reproductive organ size assessment

To test the impact of the resistance allele at the *Cyp6g1* locus and the role of nutrition and insecticide exposure on reproduction, female flies stored in fixative solution were submerged in 1 × PBS, and their ovaries removed and teased apart to count the number of ovarioles (David, 1970). Similarly, males stored in fixative solution were submerged in 1 x PBS, then one of each pair of the three reproductive organs (testis, seminal vesicle, and accessory gland) was dissected and placed on a slide in 40 µl of Dako Mounting Medium (S3023; Agilent, USA), before being sealed with a 1.0 x coverslip. Images of each structure were taken using an Olympus BX-53 microscope fitted with an Olympus 392 DP80 camera. The size of the three reproductive organs in microns was measured using ImageJ (version 1.54f).

### Statistical analysis

To test for the effect of genotype and/or insecticide exposure on each trait, models were created that include either genotype, or insecticide exposure, as fixed effects, as well as their interactions. For each addition, likelihood ratio tests between the most basic model and the newest model were performed and the most complex model was accepted if p-value<0.05. This resulted in a final full model that includes genotype, insecticide exposure, linear, and quadratic components of protein and carbohydrates, as well as a four-way interaction between genotype, insecticide exposure, protein, and carbohydrates (Supplementary Table 1).

For ovariole number no transformation was performed as the data fit a normal distribution. The construction of a model was performed similarly with stepwise additions of effects and using likelihood ratio tests to confirm the best fit model, resulting in a final model where ovariole number was used as a response variable. Genotype, insecticide exposure, linear, and quadratic components of protein and carbohydrates, as well as a four-way interaction between genotype, insecticide exposure, protein, and carbohydrates were included. Since both ovaries were dissected for each female, individual was included as an additional random effect, nested within replicate (Supplementary Table 1).

Finally, for males’ three sex organ structures, no transformations were performed as the data fit normal distributions. Similar approaches as described above were used to determine the model with the best fit. This resulted in a final model where size of each structure in micrometers squared was used as a response variable, which has been used previously as a response variable to environmental factors such as heating (Domenech & Fricke, 2022; Meena et al., 2024). Genotype, insecticide exposure, linear, and quadratic components of protein and carbohydrates, as well as a four-way interaction between genotype, insecticide exposure, protein, and carbohydrates were included as fixed effects. Since for these structures only one was measured for each individual, we only included replicate as a random effect (Supplementary Table 1).

Data was analysed and visualized in R Studio (version 3.4.1). Plots were produced by subtracting the response surfaces between corresponding variables and/or traits using ggplot2 (tidyverse package, Wickham et al., 2019). All data and R scripts are available on Figshare (to be provided).

## Results

We aimed to uncover how an insecticide resistance allele at the *Cyp6g1* locus impacts reproductive traits such as ovariole number in females, and testis, seminal vesicle, and accessory gland size in males, in response to nutrition. We furthered this knowledge by also investigating how the presence or absence of insecticides might shift the response surfaces to nutrition. All traits were measured in two genotypes (susceptible and resistant *Cyp6g1* alleles), across 25 experimental diets varying in their protein:carbohydrate ratios and caloric content, and two conditions (diets with DMSO as a vehicle control, or diets with contaminating doses of imidacloprid, an insecticide *Cyp6g1* confers resistance to).

### Effect of a *Cyp6g1* resistance allele

Diet had a significant impact on all reproductive traits observed (Figure 1, Supplementary Table 2, Supplementary Figure 1-4). These patterns revealed clear sex-specific responses to larval diet, with male and female reproductive traits differing in their macronutrient optima.

**Figure 1.**
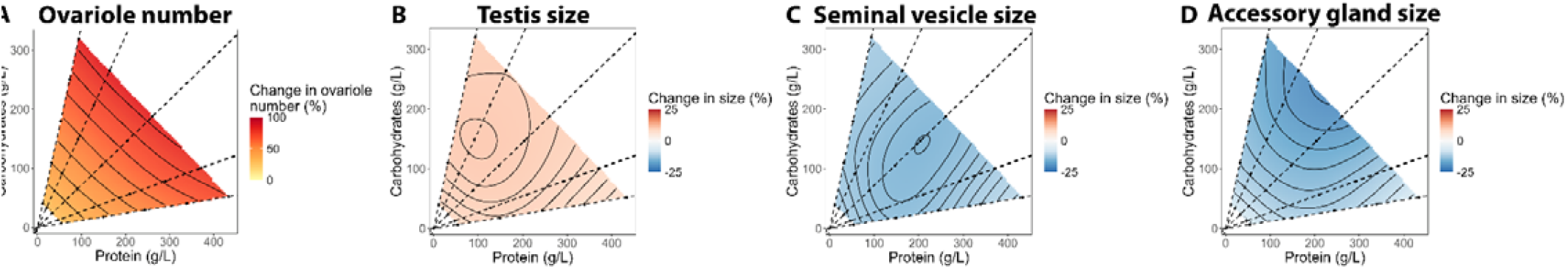
The effect of a resistance allele at the *Cyp6g1* locus on reproductive traits in response to protein and carbohydrate content of larval diet. A) Differences in ovariole number between resistant and susceptible female flies when raised on diets containing different P:C ratios and caloric contents. B-D) Differences in testis, seminal vesicle, and accessory gland size between resistant and susceptible male flies when raised on diets containing different P:C ratios and caloric contents. In all panels, dashed black lines represent protein-to-carbohydrate ratios. Each dot represents a diet on which individuals were reared. Darker red represents an increase in the trait when comparing resistant with susceptible flies, whereas darker blue colours represent a decrease in the trait when changing from a susceptible allele to a resistant allele at the *Cyp6g1* locus.

The presence of a resistance allele at the *Cyp6g1* locus also incurred significant changes in all traits observed (Figure 1, Supplementary Table 2, Supplementary Figure 1-4). Resistant female flies had an increase in ovariole number ranging between 40 to 100% compared with their susceptible counterparts (Figure 1A, Supplementary Figure 2). This difference was exacerbated with increasing caloric content in the diet (Figure 1A). For resistant males, testis size increased on average 10% across all diets tested (Figure 1B), whereas seminal vesicle and accessory gland size decreased with the presence of the resistance allele, with seminal vesicles decreasing on average by 20% across all diets (Figure 1C). Accessory glands decreased close to 25% in high caloric diets with a low protein:carbohydrate ratio, but only 5% in high protein:carbohydrate ratio diets or low caloric diets (Figure 1D).

We also observed sex-specific responses (Figure 1). Whereas both female and male gonads responded in similar way to diet and genotype with ovariole number and testis size increasing with increasing nutrition when carrying the resistance allele, this was not the case for seminal vesicle and accessory glands size. The reproductive organs of males were reduced in size when carrying the resistance allele (Figure 1). This finding highlights the interaction between *Cyp6g1*, nutrition and sex in an environment without exposure to insecticides. Furthermore, it corroborates previous findings suggesting female fitness is highest in protein-rich diets, whereas male fitness peaks in carbohydrate-rich diets (Camus et al., 2017).

### Effect of insecticide exposure in larval diet

To determine the combined effects of nutrition, genotype, and presence or absence of insecticide exposure, we measured the same reproductive traits as above in the presence of insecticides. Since insecticides are applied to food sources, even contaminating doses of insecticides are intrinsically linked to nutrition also under natural conditions.

Increasing levels of protein interacted with the presence of insecticides for ovariole number and seminal vesicle size, resulting in a reduction in size of these traits (Figure 2, Supplementary Table 2). For ovariole number this effect was exacerbated by the genotype effect, with susceptible flies having a greater reduction in ovariole number with increasing protein in the diet and in the presence of insecticides (Figure 2A, Supplementary Table 2). Moreover, a 4-way interaction was detected between genotype, protein, carbohydrates and insecticide exposure for ovariole number and accessory gland size (Supplementary Table 2). This response was evident by an increase in ovariole number and accessory gland size in the presence of insecticides, but only within a certain dietary space that varied depending on the *Cyp6g1* allele present (Figure 2A, D, Supplementary Table 2). Insecticide exposure also significantly impacted all other male traits independently, resulting in an increase in testis, and seminal vesicle size (Figure 2B, C, Supplementary Table 2, Supplementary Figures 2-4).

**Figure 2.**
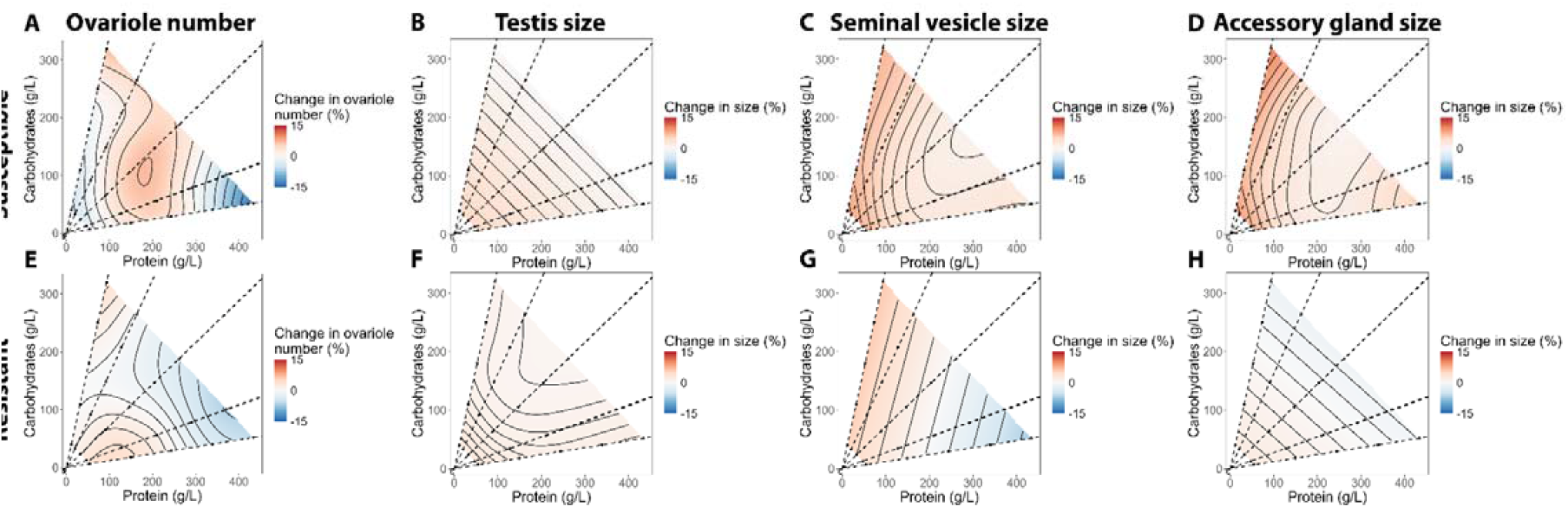
The effect of insecticide exposure on reproductive trait size in response to protein and carbohydrate content of larval diet. A-D) Differences in ovariole number, testis, seminal vesicle, and accessory gland size of susceptible *Drosophila melanogaster* when exposed to nsecticides. E-H) Differences in ovariole number, testis, seminal vesicle, and accessory gland size of resistant *Drosophila melanogaster* when exposed to insecticides. In all panels, dashed black lines represent protein-to-carbohydrate ratios. Each dot represents a diet on which ndividuals were reared. Darker red represents an increase of the trait when adding insecticides to the diet, whereas darker blue represents a decrease in the trait when adding insecticides to the diet.

Beyond the effects of diet and genotype described above, insecticide exposure significantly altered how fertility traits responded to larval nutrition. The effects of diet on reproductive traits were not fixed but shifted in the presence of insecticides. This effect was reflected by multiple significant interaction terms between diet, genotype, and insecticide exposure. These interactions demonstrate that insecticides do not act as independent stressors but are intrinsically embedded within the nutritional environment experienced by insects. Our findings revealed that reproductive trait size emerges from the interaction between resistance alleles, insecticide exposure, and diet, exposing a critical and previously absent interaction in insecticide research.

## Discussion

In this study, we aimed to understand how insecticide resistance alleles shape nutritional performance of each sex in the presence or absence of insecticides. We studied nutritional performance and insecticide exposure in two *Cyp6g1* alleles of *D. melanogaster* that confer resistance or susceptibility to a wide range of insecticides. Given the difference in resistance and the different nutritional requirements of various reproductive traits, we hypothesized that resistant individuals would outperform susceptible flies in environments with insecticides, but not always in environments without this toxin. Furthermore, previous studies have shown that the resistant allele in the *Cyp6g1* locus also provides a boost in fecundity in female flies (McCart et al., 2005; Rostant et al 2015). However, this fecundity enhancement was only examined in one diet, and so we expect this increase in fecundity to fluctuate depending on the nutritional context the larvae are reared in.

Our results demonstrate that nutrition is a critical regulator of reproductive trait size, while also revealing that its effects are fundamentally reshaped by insecticide resistance alleles, sex, and insecticide exposure. Consistent with extensive prior work, larval diet exerted a strong influence on fertility-related traits, with protein-rich diets enhancing female ovariole number (Lee et al., 2008; Mirth et al., 2019; Rodrigues et al., 2015). These findings reinforce the central role of nutrition in shaping life-history traits. However, our study extends this framework by showing that male reproductive traits such as testis, seminal vesicle, and accessory gland sizes are also highly sensitive to diet, genotype, and their interaction – a dimension that to date has been comparatively understudied.

A key finding is that the fitness consequences of the *Cyp6g1* resistance allele are strongly sex-and diet-dependent. Resistant females consistently showed increased ovariole number regardless of diet, confirming previous observations that resistance alleles can enhance female fecundity (McCart et al., 2005; Rostant et al 2015). Importantly, this benefit was amplified under high-calorie and protein-rich diets, suggesting that resistance-associated reproductive gains are maximised in a dietary environment different from the one that optimises fecundity of susceptible females. In contrast, resistant males exhibited a more complex pattern: although testis size increased modestly, seminal vesicles and accessory gland sizes were substantially reduced, particularly under carbohydrate-rich, high-calorie diets. Given the central role of these organs in sperm storage, seminal fluid production, and post-mating female responses, such reductions may translate into reduced male reproductive performance (Bangham et al., 2002; Santhosh & Krishna, 2013). Although further studies will be needed to determine if changes in accessory gland size modify ejaculate composition between susceptible and resistant male flies, previous studies show that accessory gland size determines the reproductive success of males (Bangham et al., 2002). Furthermore, bigger accessory glands are also correlated with higher transfer of accessory gland proteins (ACPs) to the female through the ejaculate (Wigby et al., 2009). Together, these results highlight that resistance evolution can reshape reproductive trade-offs differently in males and females. Moreover, they reinforce the idea that nutritional optima and fitness consequences cannot be generalised across sexes, especially when taking account *Cyp6g1* resistance alleles.

Insecticide exposure further altered the dynamics between susceptible and resistant flies in unexpected ways. Contaminating doses of imidacloprid reshaped the nutritional landscape, modifying in which diets the reproductive traits would reach an optimum. Notably, in susceptible flies, low-level insecticide exposure increased the size of male reproductive organs and, within specific dietary spaces, enhanced ovariole number in females. This phenomenon indicates the possibility of hormesis regulating the response to nutrition and insecticide exposure (Mattson, 2008). Hormesis is a phenomenon where a small environmental stress protects against future non-lethal stresses (Mattson, 2008). In this case, the presence of low doses of insecticides might trigger stress-responses that protect the larvae from further nutritional stress, or inversely, nutritional stress might protect against low doses of insecticides, enabling them to allocate resources to develop bigger sexual organs. This stimulatory effect was most evident in susceptible genotypes, whereas resistant flies failed to derive comparable benefits. One plausible explanation is that resistant individuals may already experience the maximum benefit due to sustained overexpression or activity of *Cyp6g1*. In the presence of insecticides, *Cyp6g1*’s detoxification machinery may be preferentially recruited to detoxify the insecticide and reduce its interaction with nutrition, constraining resource allocation to reproduction when insecticides are present, even at contaminating levels.

Overall, our results demonstrate that resistance alleles play more roles than simply enabling survival under chemical stress; they actively reshape reproductive allocation across nutritional landscapes. By demonstrating that insecticide exposure also reshapes nutritional optima in a sex-specific manner, we reveal that resistance evolution is embedded within dietary landscapes rather than driven by toxicity alone. In natural and agricultural systems, insecticides are inseparable from food resources, yet resistance is typically studied in simplified contexts that ignore this ecological reality. Our findings highlight that shifts in allele frequency, reproductive output, and population persistence will depend on how toxin stress interacts with nutrient availability. Integrating nutritional ecology into the study of resistance is therefore essential for predicting evolutionary trajectories in human-modified environments.

## Supplementary Figures and Tables

**Supplementary Figure 1.**
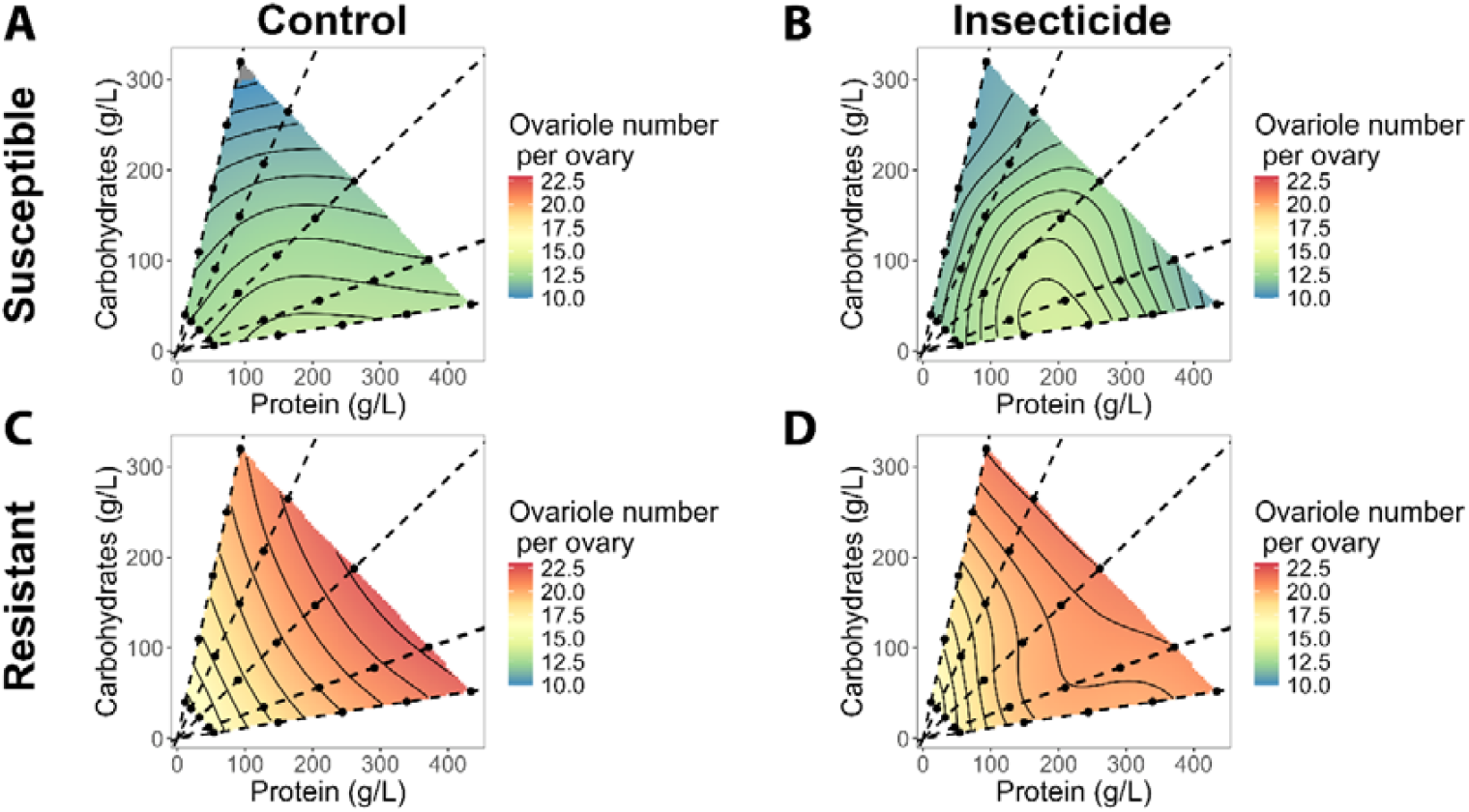
Changes in ovariole number in response to protein and carbohydrate content of larval diet for two different alleles of *Cyp6g1* (A, B, susceptible allele; C, D, resistant allele) and in the presence or absence of a contaminating dose of imidacloprid (A, C, control food dosed with DMSO; B, D, food dosed with imidacloprid). In all panels dashed black lines represent protein-to-carbohydrate ratios. Each dot represents a diet on which individuals were reared.

**Supplementary Figure 2.**
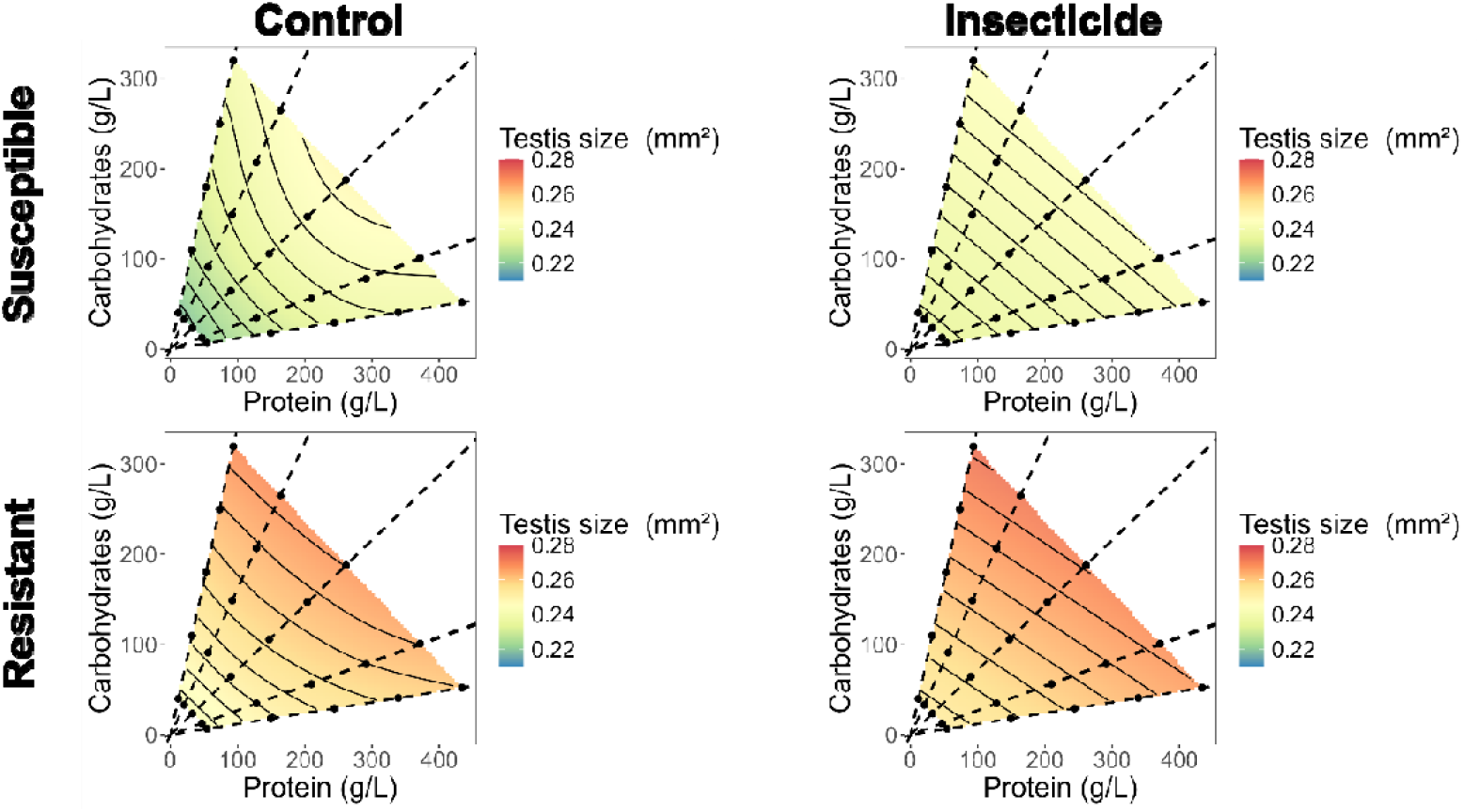
Changes in testis size in response to protein and carbohydrate content of larval diet for two different alleles of *Cyp6g1* (A, B, susceptible allele; C, D, resistant allele) and in the presence or absence of a contaminating dose of imidacloprid (A, C, control food dosed with DMSO; B, D, food dosed with imidacloprid). In all panels dashed black lines represent protein-to-carbohydrate ratios. Each dot represents a diet on which individuals were reared.

**Supplementary Figure 3.**
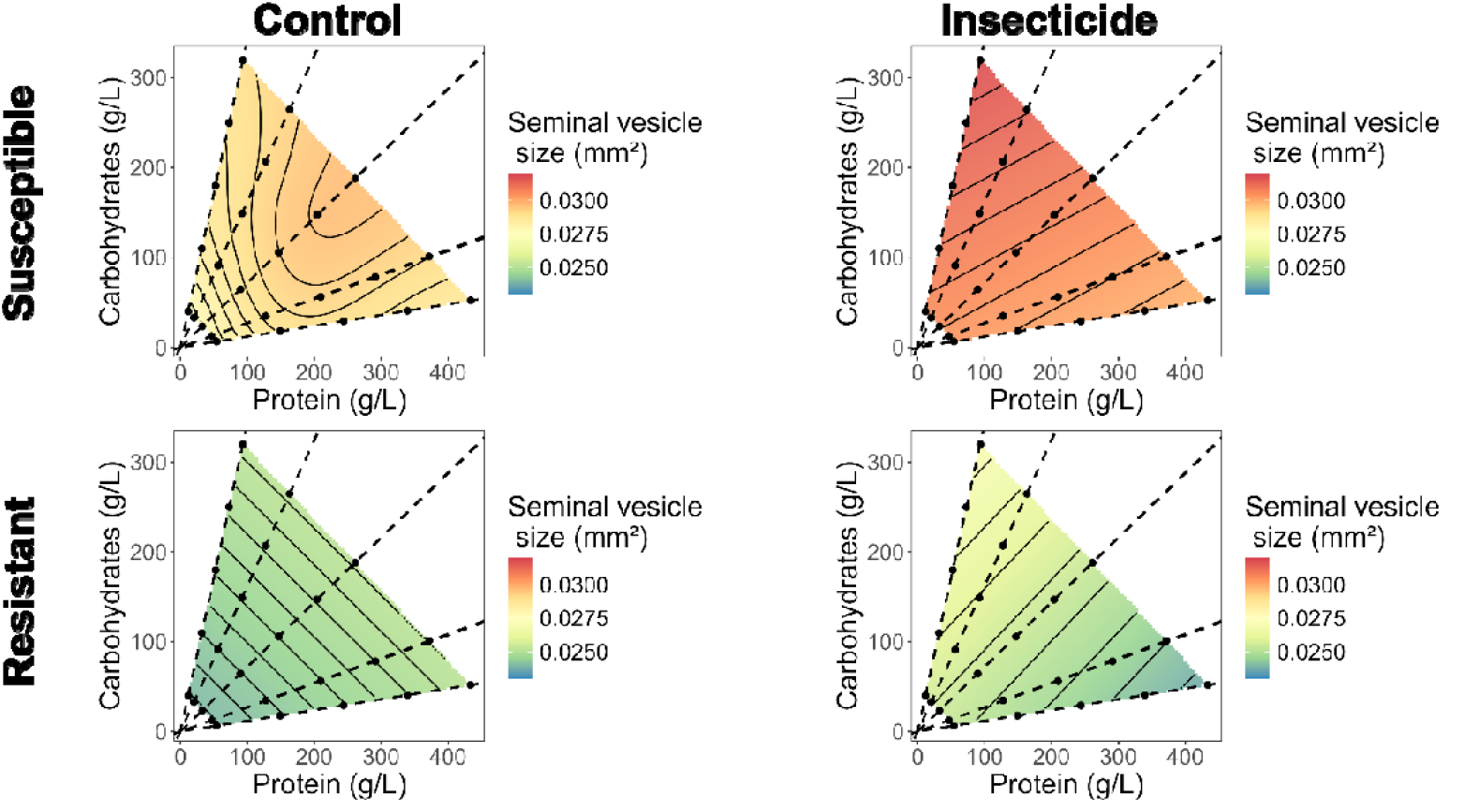
Changes in seminal vesicle size in response to protein and carbohydrate content of larval diet for two different *Cyp6g1* alleles (A, B, susceptible allele; C, D, resistant allele) and in the presence or absence of a contaminating dose of imidacloprid (A, C, control food dosed with DMSO; B, D, food dosed with imidacloprid). In all panels dashed black lines represent protein-to-carbohydrate ratios. Each dot represents a diet on which individuals were reared.

**Supplementary Figure 4.**
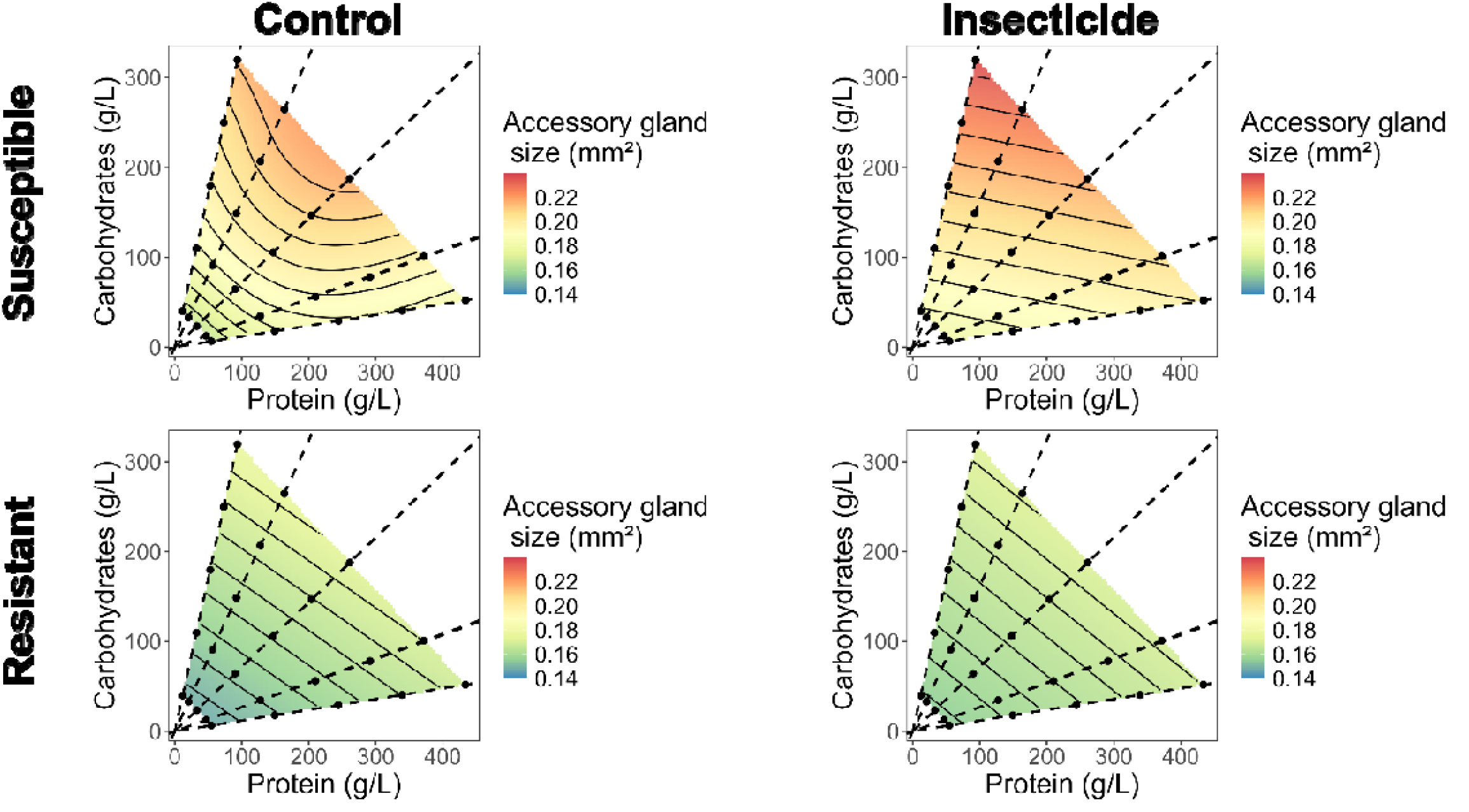
Changes in accessory gland size in response to protein and carbohydrate content of larval diet in males with two different *Cyp6g1* alleles (A, B, susceptible allele; C, D, resistant allele) and in the presence or absence of a contaminating dose of imidacloprid (A, C, control food dosed with DMSO; B, D, food dosed with imidacloprid). In all panels dashed black lines represent protein-to-carbohydrate ratios. Each dot represents a diet on which individuals were reared.

**Supplementary Table 1:**
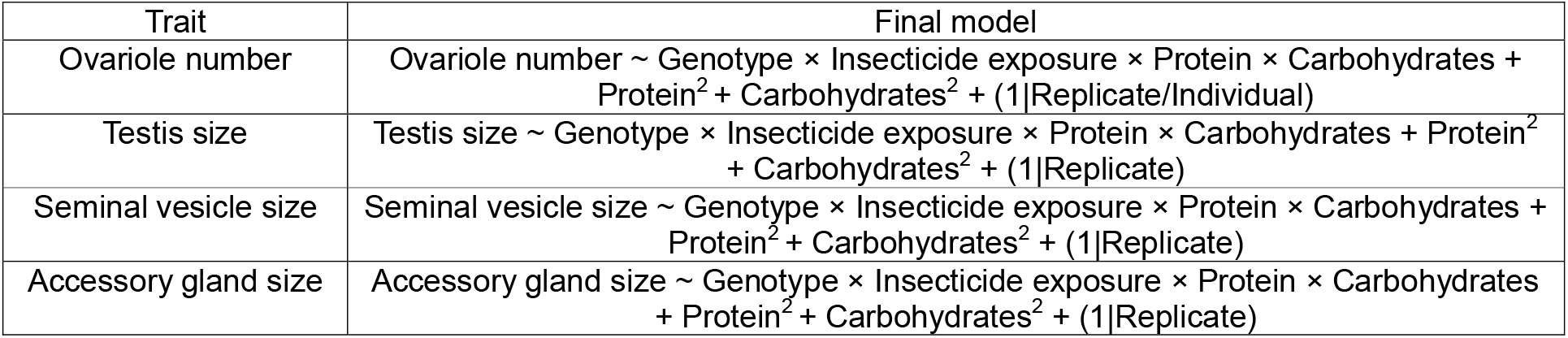
Final linear mixed effects models used to analyse the effect of genotype, insecticide exposure, and nutrition on life-history traits.

**Supplementary Table 1:** The effect of genotype, insecticide presence, carbohydrates, protein and their interactions on development time, ovariole number, testis, seminal vesicle, and accessory gland size. Data were analysed using linear models with mixed effects. Significant terms in bold.

| Term | Ovariole number |  |  | Testis size |  |  | Seminal vesicle size |  |  | Accessory gland size |  |  |
| --- | --- | --- | --- | --- | --- | --- | --- | --- | --- | --- | --- | --- |
|  | Chisq | DF | p-value | Chisq | DF | p-value | Chisq | DF | p-value | Chisq | DF | p-value |
| Genotype | 3636.35 | 1 | **<2.2e-16 \*\*\*** | 329.93 | 1 | **<2.2e-16 \*\*\*** | 231.5 | 1 | **<2.2e-16 \*\*\*** | 413.73 | 1 | **<2.2e-16 \*\*\*** |
| Insecticide exposure | 0.04 | 1 | 0.846 | 18.4 | 1 | **1.79e-05 \*\*\*** | 7.46 | 1 | **0.0063 \*\*** | 8.66 | 1 | **0.0033 \*\*** |
| Carbohydrates | 0.4 | 1 | 0.5276 | 17.95 | 1 | **2.27e-05 \*\*\*** | 0.02 | 1 | 0.9007 | 31.55 | 1 | **1.95e-08 \*\*\*** |
| Protein | 292.46 | 1 | **<2.2e-16 \*\*\*** | 14.52 | 1 | **0.0001 \*\*\*** | 7.69 | 1 | **0.0055 \*\*** | 43.99 | 1 | **3.30e-11 \*\*\*** |
| I(Carbohydrates^2) | 0.92 | 1 | 0.3385 | 3.8 | 1 | 0.0513 | 0.14 | 1 | 0.7093 | 8.62 | 1 | **0.0033 \*\*** |
| I(Protein^2) | 172.28 | 1 | **<2.2e-16 \*\*\*** | 4.41 | 1 | **0.0358 \*** | 7.81 | 1 | **0.0052 \*\*** | 26.33 | 1 | **2.87e-07 \*\*\*** |
| Genotype * Insecticide exposure | 0.2 | 1 | 0.6573 | 0.5 | 1 | 0.4815 | 3.41 | 1 | 0.0648 | 8.77 | 1 | **0.0031 \*\*** |
| Genotype * Carbohydrates | 154.78 | 1 | **<2.2e-16 \*\*\*** | 1.97 | 1 | 0.161 | 0.01 | 1 | 0.9182 | 24.4 | 1 | **7.83e-07 \*\*\*** |
| Insecticide exposure * Carbohydrates | 0.05 | 1 | 0.8271 | 3.43 | 1 | 0.0641 | 0.29 | 1 | 0.5913 | 0.04 | 1 | 0.8439 |
| Genotype * Protein | 123.2 | 1 | **<2.2e-16 \*\*\*** | 0.07 | 1 | 0.7871 | 0.6 | 1 | 0.4386 | 0.02 | 1 | 0.884 |
| Insecticide exposure * Protein | 15.32 | 1 | **9.06e-05 \*\*\*** | 1.75 | 1 | 0.1855 | 6 | 1 | **0.0143 \*** | 2.51 | 1 | 0.1131 |
| Carbohydrates * Protein | 5.71 | 1 | **0.0169 \*** | 0.06 | 1 | 0.8124 | 0.37 | 1 | 0.5411 | 0.42 | 1 | 0.5163 |
| Genotype * Insecticide exposure * Carbohydrates | 1.14 | 1 | 0.2848 | 0.34 | 1 | 0.5577 | 0.19 | 1 | 0.6641 | 0.27 | 1 | 0.606 |
| Genotype * Insecticide exposure * Protein | 4.51 | 1 | **0.0337 \*** | 0.53 | 1 | 0.4669 | 0.97 | 1 | 0.3245 | 0.01 | 1 | 0.9263 |
| Genotype * Carbohydrates * Protein | 1.38 | 1 | 0.2408 | 0.4 | 1 | 0.5256 | 4.81 | 1 | **0.0283 \*** | 0.22 | 1 | 0.6403 |
| Insecticide exposure * Carbohydrates * Protein | 0.76 | 1 | 0.3828 | 0.15 | 1 | 0.6984 | 0.58 | 1 | 0.4462 | 0.55 | 1 | 0.458 |
| Genotype * Insecticide exposure * Carbohydrates * Protein | 4.53 | 1 | **0.0334 \*** | 0.02 | 1 | 0.8929 | 1.37 | 1 | 0.2419 | 5.79 | 1 | **0.0161 \*** |

